# Trends in CRP, D-dimer and fibrinogen during therapy for HIV associated multidrug resistant tuberculosis

**DOI:** 10.1101/268367

**Authors:** Patrick G. T. Cudahy, Joshua L. Warren, Ted Cohen, Doug Wilson

## Abstract

**Background:** HIV positive adults on treatment for multidrug-resistant tuberculosis (MDR-TB) experience high mortality. Biomarkers of HIV/MDR-TB treatment response may enable earlier treatment modifications that improve outcomes.

**Methods:** To determine whether trends in C-reactive protein (CRP), D-dimer and fibrinogen predict treatment outcome among those with HIV/MDR-TB co-infection we studied 20 HIV positive participants initiating therapy for MDR-TB. Serum CRP, fibrinogen, and D-dimer were measured at baseline and serially while on treatment. Results: At baseline, all biomarkers were elevated with median CRP 86.15 mg/L (IQR 29.25-149.32), D-dimer 0.85 μg/mL (IQR 0.34-1.80) and fibrinogen 4.11 g/L (IQR 3.75-6.31). CRP decreased significantly within 10 days of treatment initiation and fibrinogen within 28 days; D-dimer did not change significantly. 5 (25%) participants died. Older age (median age of 38y among survivors and 54y among deceased, p=0.008) and higher baseline fibrinogen (3.86 g/L among survivors and 6.37 g/L among deceased, p=0.02) were significantly associated with death. Higher CRP concentrations at the beginning of each measurement interval were significantly associated with a higher risk of death during that interval.

**Conclusion:** Trends in fibrinogen and CRP may be useful for evaluating early response to treatment among individuals with HIV/MDR-TB co-infection.

## INTRODUCTION

Tuberculosis (TB) remains the leading worldwide infectious cause of death^1^, driven in part by high mortality in HIV co-infected adults and by the emergence of strains of *M. tuberculosis* resistant to both isoniazid and rifampicin (multidrug resistant tuberculosis [MDR-TB]). New diagnostics have been broadly deployed but significant population level improvements in outcomes have not followed^2^. Especially concerning are the high levels of early mortality (within 60 days) experienced by patients beginning therapy for MDR-TB in South Africa, most of whom are HIV co-infected^3^.

New molecular diagnostics have improved the ability to rapidly identify the presence of drug-resistance but often provide only limited information about susceptibility profiles of the infecting strain of *M. tuberculosis*. The Xpert MTB/Rif, currently the most widely used rapid molecular test, provides information on rifampicin resistance only^4^. Direct line-probe assays on sputum are also utilized for detecting first and second line drug resistance but have low sensitivity in smear negative disease^5^, which is common in HIV associated tuberculosis^6^. More typically, several weeks to months are required before phenotypic drug susceptibility testing (DST) or indirect line-probe assays are available to guide therapy. Therefore, standardized second-line regimens are used during the period between diagnosis and confirmation of drug susceptibility by culture dependent methods, when risk of death is greatest^7^.

There are currently no clinically useful early markers of effective therapy^8^. The earliest World Health Organization (WHO) recommended monitoring is sputum smear microscopy for acid-fast bacilli at the completion of two months of treatment^9^. However, mortality at two months is already very high in certain populations and settings^7^ and many HIV co-infected individuals are smear-negative at the time of TB diagnosis^10^, limiting the effectiveness of smear monitoring among these co-infected patients at highest risk of death. Earlier biomarkers of treatment response could identify individuals not responding rapidly to therapy, allow for earlier modification of ineffective treatment and potentially prevent deaths.

Such biomarkers would be most valuable if they could be immediately used by clinicians at patient bedsides in high burden, low resource settings. We therefore focused on candidate markers of early MDR-TB treatment response which were available in routine clinical practice in South Africa, and available on point-of-care platforms in the developed world^11^.

C-reactive protein (CRP) is elevated in 93% of patients with HIV and TB co-infection at time of diagnosis^12^. Once on therapy, studies of individuals without HIV-coinfection and with drug-sensitive TB have shown reductions in CRP levels to be associated with increased two month culture conversion rates and less need for treatment extension^13–16^. Fibrinogen has also been shown to be elevated in HIV-TB co-infected patients^17,18^ and to normalize within the first two months of therapy^19,20^. Elevated D-dimer has been associated with pulmonary tuberculosis^17^ and HIV/TB co-infection^21^. We conducted a study to prospectively evaluate these three candidate markers of treatment response among people living with HIV beginning treatment for MDR-TB.

## METHODS

Doris Goodwin Hospital (DGH) is a dedicated public MDR-TB treatment facility in KwaZulu-Natal, South Africa. It serves as the primary MDR-TB treatment site of the uMgungundlovu Health District. In 2015, the district had a case notification rate for tuberculosis of 678 per 100,000; 6.4% of all tuberculosis patients had rifampin resistance and 70.2% were HIV co-infected^22^.

We enrolled twenty consecutive HIV-positive patients admitted to DGH for initiation of MDR-TB therapy between September 2016 and March 2017. Inclusion criteria were: 1) age greater than or equal to 18 years; 2) sputum positive for *M. tuberculosis* with rifampin resistance by molecular or phenotypic drug susceptibility testing; 3) HIV seropositive; 4) able to attend clinic visits at DGH after discharge; 5) willing to give informed consent for participation. Exclusion criteria were: 1) known resistance at baseline to fluoroquinolones or kanamycin/amikacin/capreomycin; 2) initiation of HIV antiretroviral therapy within 12 weeks prior to initiating MDR-TB treatment; 3) pregnancy; 4) prisoners.

After the informed consent process, venous blood was collected at baseline before treatment initiation, and at days 5, 10, 14 and 28, and weeks 8, 12 and 16 after treatment initiation, either while hospitalized or during regular clinic follow-up appointments. Blood was collected in vacuum blood collection tubes, stored at ambient temperature and tested within 8 hours of collection. Samples were supplied to laboratory personnel with a coded participant identifier. Laboratory personnel were blinded to all other participant information. CRP was determined by immunoturbidimetric assay (Siemens Healthcare (Pty) Ltd, S.A, normal range 0 to 5 mg/L), D-dimer was determined by reflectometric quantitative measurement (Roche Products (Pty) Ltd, S.A, normal range 0.1 to 0.5 μg/mL) and fibrinogen by immunologic method (Sysmex South Africa (Pty) Ltd, S.A, normal range 2 to 4 g/L), all according to manufacturer protocols. Thresholds for the upper limit of normal were set by the respective manufacturers. Hemoglobin, creatinine, albumin and alkaline phosphatase results were obtained by review of participant records after tests were routinely performed by the South African National Health Laboratory Service.

Participant records were examined to determine demographic and additional medical information. Tuberculosis treatment was conducted according to local guidelines by department of health clinicians. Clinical staff were blinded to the results of the research assays. At the time of the study, MDR-TB patients were initiated on a standardized regimen, including moxifloxacin, kanamycin, and at least 3 additional agents^23^. Those with severe baseline hearing loss or renal impairment started a bedaquiline-based regimen, and those developing severe side-effects on fluoroquinolones or injectables were switched to a bedaquiline-based regimen, per local guidelines.

Participants not on HIV antiretroviral therapy (ART) at baseline were started on MDR-TB therapy first, and subsequently fast-tracked onto ART, per South African guidelines^24^.

### Statistical Analysis

Comparisons between groups for continuous variables were tested with Kruskal-Wallis and for categorical variables by chi-square tests. A discrete time survival model^25,26^ was used to describe the hazard of death during each time interval controlling for age category of the patient (greater than or equal to 50 vs. less than 50 years), changing risk across time (linear and quadratic time effects), and temporally varying biomarker measurements for the patient. The model was fit using a binary variable (survival/death during each time interval for each patient) regression approach with the complementary log-log link function in R statistical software^27^. Cutoffs for continuous biomarkers were determined by the Youden index of the receiver operating characteristic (ROC) curve. Laboratory clinical severity index was calculated as previously described^28^. A score from 0 to 4 was assigned with one point for each abnormal value in hemoglobin, creatinine, albumin, or alkaline phosphatase, with cutoffs defined by our laboratory’s standards, with hemoglobin of < 13g/L, creatinine > 104 μmol/L, albumin < 35 g/L and alkaline phosphatase > 128 IU/L considered abnormal.

Data were collected using REDCap electronic data capture tools hosted at Stanford University^29^, and analyzed using jupyter version 4.4.0 (jupyter.org) and R version 3.4.1 (r-project.org)^27^ and with packages tableone version 0.8.1, ggplot2 version 2.2.1^30^, and pROC version 1.10.0^31^.

### Ethics

Written consent was obtained. The study was approved by the South African Medical Association Research Ethics Committee and Yale University Human Investigation Committee. All research was performed in accordance with relevant guidelines and regulations.

### Data availability

The datasets generated during and/or analyzed during the current study are available from the corresponding author on reasonable request.

## RESULTS

### Study Population

98 potential participants were screened for entry into the study. Of these, 64 were ineligible (24 started TB therapy before they could be approached for consent, 17 were HIV negative, 9 were transferred in to the facility already on TB treatment, 5 were already enrolled in another study, 5 were prisoners, 2 had been started on antiretroviral therapy in the 12 weeks before admission, and 2 were unable to provide informed consent due to altered mental state). 36 individuals were approached for enrollment and 20 consented to participate. None of the participants withdrew consent at a later date and all were followed for at least 16 weeks or until death.

Most participants were male (n=13), with median age of 41 years (inter-quartile range (IQR) 33 to 48y) (Table 1). 12 had been treated previously for tuberculosis and 2 of these individuals had prematurely stopped their prior course of treatment. The median CD4 cell count was 234 cells per μL (IQR of 141 to 429 cells/μL), 12 were on antiretroviral therapy (ART), and 10 of these individuals had a recent undetectable viral load measurement. Those not on ART at baseline were initiated at a median of 12.5 days after starting TB therapy (range 7 to 86 days). Participants were admitted to an inpatient unit at the initiation of therapy for a median of 32 days (range 5 to 140 days).

**Table 1.** Participant baseline characteristics

|  | Category | Value | Normal range |
| --- | --- | --- | --- |
| n |  | 20 |  |
| Sex (%) | Female | 7 (35.0) |  |
|  | Male | 13 (65.0) |  |
| Race (%) | Indian | 1 (5.0) |  |
|  | Black African | 19 (95.0) |  |
| Age (years) (median [IQR]) |  | 41.95 [33.76, 48.93] |  |
| BMI* (median [IQR]) |  | 19.35 [16.23, 24.67] |  |
| ECOG<sup>†</sup> (%) | 1 | 13 (65.0) |  |
|  | 2 | 4 (20.0) |  |
|  | 3 | 3 (15.0) |  |
| TB category (%) | New TB case | 8 (40.0) |  |
|  | Prior history of TB | 10 (50.0) |  |
|  | Treatment after default | 2 (10.0) |  |
| Sputum smear grade (%) | 0 | 12 (60.0) |  |
|  | 1 | 1 (5.0) |  |
|  | 2 | 3 (15.0) |  |
|  | 4 | 4 (20.0) |  |
| CD4 count (cells/μL) (median [IQR]) | | 234.00 [141.75, 429.75] | |
| Detectable viral load (%) | Detectable viral load on ART<sup>‡</sup> | 2 (10.0) |  |
|  | Undetectable viral load on ART | 10 (50.0) |  |
|  | Not on ART | 8 (40.0) |  |
| Hemoglobin (g/L) (median [IQR]) |  | 10.05 [9.38, 11.33] | 13.0-17.0 |
| Creatinine (μmol/L) (median [IQR]) | | 72.50 [68.00, 85.25] | 64-104 |
| Albumin (g/dL) (median [IQR]) |  | 24.00 [21.00, 28.50] | 35-52 |
| Alkaline phosphatase (IU/L) (median [IQR]) |  | 125.00 [111.25, 145.00] | 53-128 |
| LCSI<sup>§</sup> (%) | 2 | 11 (55.0) |  |
|  | 3 | 7 (35.0) |  |
|  | 4 | 2 (10.0) |  |
| CRP (mg/L) (median [IQR]) |  | 86.15 [29.25, 149.32] | 0.0-5.0 |
| Fibrinogen (g/L) (median [IQR]) |  | 4.11 [3.75, 6.31] | 2.00-4.00 |

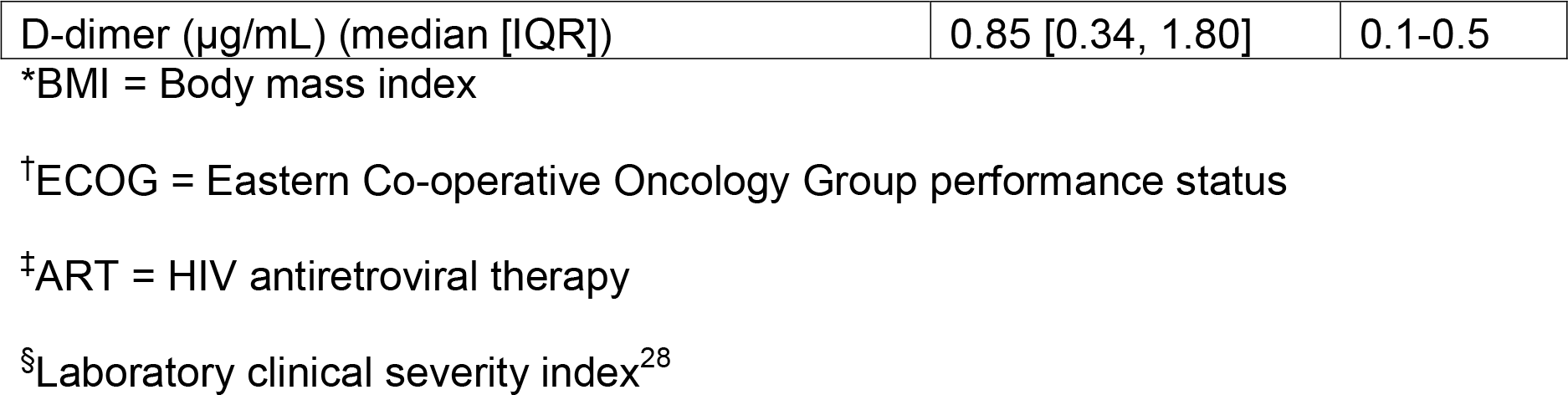

At baseline, almost all participants were anemic and hypoalbuminemic; 45% had an elevated alkaline phosphatase and 15% had baseline renal impairment. CRP, fibrinogen and D-dimer were all elevated at baseline with a median CRP of 86.15mg/L (upper limit of normal (ULN) of 5mg/L, IQR 29.25 to 149.32 mg/L), median fibrinogen of 4.11 g/L (ULN of 4 g/L, IQR 3.75 to 6.31 g/L), and median D-dimer of 0.85 μg/mL (ULN 0.5 μg/mL IQR 0.34 to 1.80 μg/mL).

### Trends

While on therapy for MDR-TB, median CRP and fibrinogen levels decreased significantly while D-dimer did not (Supplementary Figure 1). CRP fell significantly by day 10 of therapy, and fibrinogen by day 28. The reductions were sustained beyond day 28, with median fibrinogen (but not CRP) reaching normal range.

### Outcomes

After four months of follow-up, 5 of the 20 participants died (25%) at a median of 32 days after starting TB treatment (range 4 to 51 days).

At baseline, older age and higher baseline fibrinogen level were significantly associated with higher risk of death. Median age was 54 years (IQR 48 to 60y) among those dying during treatment and 38 years (IQR 32 to 44y) among survivors, p=0.008. Median baseline fibrinogen was 6.37 g/L (IQR 6.05 to 7.50 g/L) for those dying and 3.86 g/L (IQR 3.68 to 4.33 g/L) for survivors, p=0.02. Baseline CRP was elevated among those that died with a median of 117 mg/L (IQR 87.9 to 169.7 mg/L), but this was not statistically different than the baseline CRP among survivors (median 76.6 mg/L, IQR 17.8 to145.7 mg/L), p=0.127. Baseline D-dimer was similar in both survivors and non-survivors (Figure 1).

**Figure 1.**
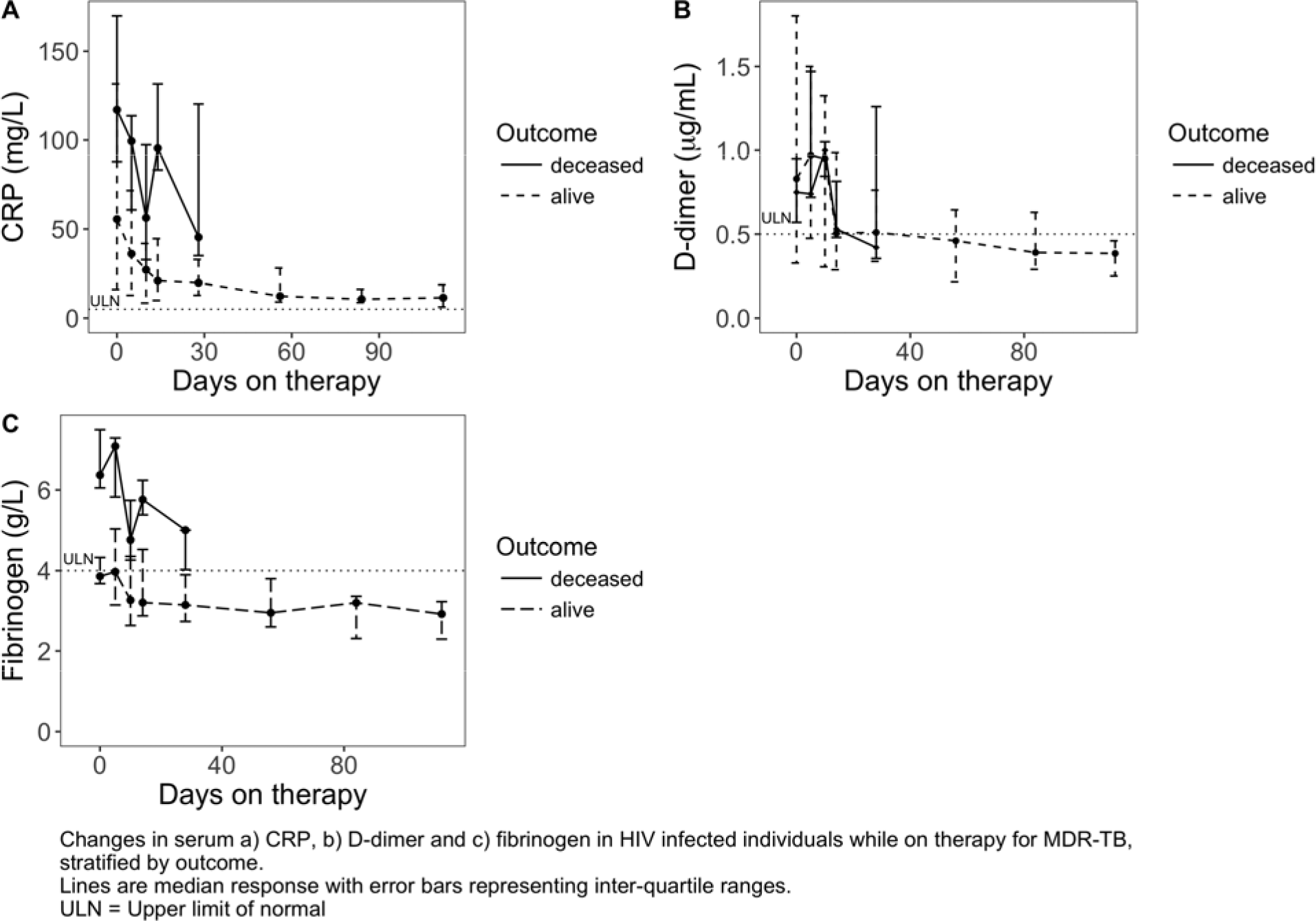
Outcome group trends while on MDR-TB therapy

Hemoglobin, creatinine, albumin, and alkaline phosphatase were not significantly different at baseline between survivors and non-survivors.

Results from the discrete time survival model suggest that patients with higher CRP values at the beginning of a measurement interval had a significantly higher risk of dying during that time interval. All other factors being equal for two living patients at the start of a time interval, a patient whose CRP value is higher (by the interquartile range of CRP observed across all patients) had an estimated increase in the hazard of dying during that time interval of 194% (95% CI: 25% to 591%).

The discrete time survival model did not show statistically significant associations between patients with higher fibrinogen or d-dimer at time interval start and hazard of dying during the time interval. For fibrinogen, larger interquartile values had an estimated increase in the hazard of dying by 171.08% (95% CI: −51.25% to 1407.23 and for D-dimer by 126.01% (95% CI: −34.29% to 677.39%).

Failure to reduce CRP by 44% at week 2 had a sensitivity of 71% and specificity of 100% for predicting death (ROC area under the curve of 0.74, 95% CI 0.50 to 0.97).

## DISCUSSION

We conducted a study that prospectively followed a cohort of 20 participants with HIV and MDR-TB co-infection for the first 4 months of their therapy and found that individuals who had higher CRP concentrations at the start of a time interval had a significantly increased hazard of dying. The relative decrease in CRP in survivors was similar to that seen in other work by our group in people with HIV and drug-sensitive TB (manuscript in preparation) with a failure to reduce CRP by 44% at week 2 the most sensitive cutoff of predicting death. Baseline fibrinogen was also significantly higher in those that died.

Older age was a significant baseline predictor of death, a finding which has been reported by others in South Africa^32^. Other routinely collected clinical parameters were not significantly predictive of outcome.

Median CRP and fibrinogen rapidly declined on MDR-TB therapy, with CRP having a significant decrease by 10 days, and fibrinogen by 28 days on treatment. The persistently elevated fibrinogen and CRP in those who died could have been a result of ineffective TB treatment, an unfavorable inflammatory response phenotype, coinfection with another opportunistic disease, advanced HIV disease, or unrelated medical illness. Our results suggest that failure to normalize CRP and fibrinogen may be useful signals to prompt clinicians to risk stratify patients on therapy with the intention of pursuing further investigations or altering therapy. Earlier effective therapy may improve outcomes as well as reduce the risk of generating additional resistance mutations and halt further drug resistant TB transmission^33^.

D-dimer was elevated at time of diagnosis for most participants but was not predictive of outcome. During the course of therapy there was no significant difference between D-dimer trends in survivors and deaths and overall, average D-dimer levels did not decline significantly.

Other routinely collected laboratory values (hemoglobin, creatinine, albumin, and alkaline phosphatase) measured at baseline were similar between the two groups, though abnormal in almost all participants. While studying patients with XDR-TB in the same setting, Shenoi et al. developed a laboratory clinical severity index (LCSI)^28^. In our population this score was not significantly predictive of outcome, most likely because all participants had baseline derangements in at least two of these four values, and 45% had derangements in at least three, leaving little room for comparison.

Strengths of our study are that data was obtained by sequential recruitment of participants with confirmed rifampin-resistant TB at a public treatment site, who were treated with standardized regimens according to national guidelines. Treating physicians were blinded to CRP, D-dimer and fibrinogen results. We note that this was a small study and the results will need to be followed up within larger cohorts. Furthermore, 40% of our study’s participants were not taking ART at baseline, and initiation of ART that occurred during MDR-TB treatment may have triggered immune reconstitution, complicating interpretation of observed biomarker trends.

## CONCLUSION

Our analysis of CRP and fibrinogen trends suggests that these biomarkers may predict mortality in HIV positive adults on treatment for MDR-TB and should be further evaluated in larger studies.

## Acknowledgements

We would like to thank our study nurses, S Mbutho and L Maziya; the KwaZulu-Natal Department of Health; and the clinical staff and management at Doris Goodwin Hospital for their commitment to improving tuberculosis care.

## Contributions

PGTC, TC and DW conceived of and designed the study. PGTC collected the data. PGTC and JLW analyzed the data. PGTC, TC, JLW and DW wrote the manuscript.

---

**Supplementary Figure 1:**
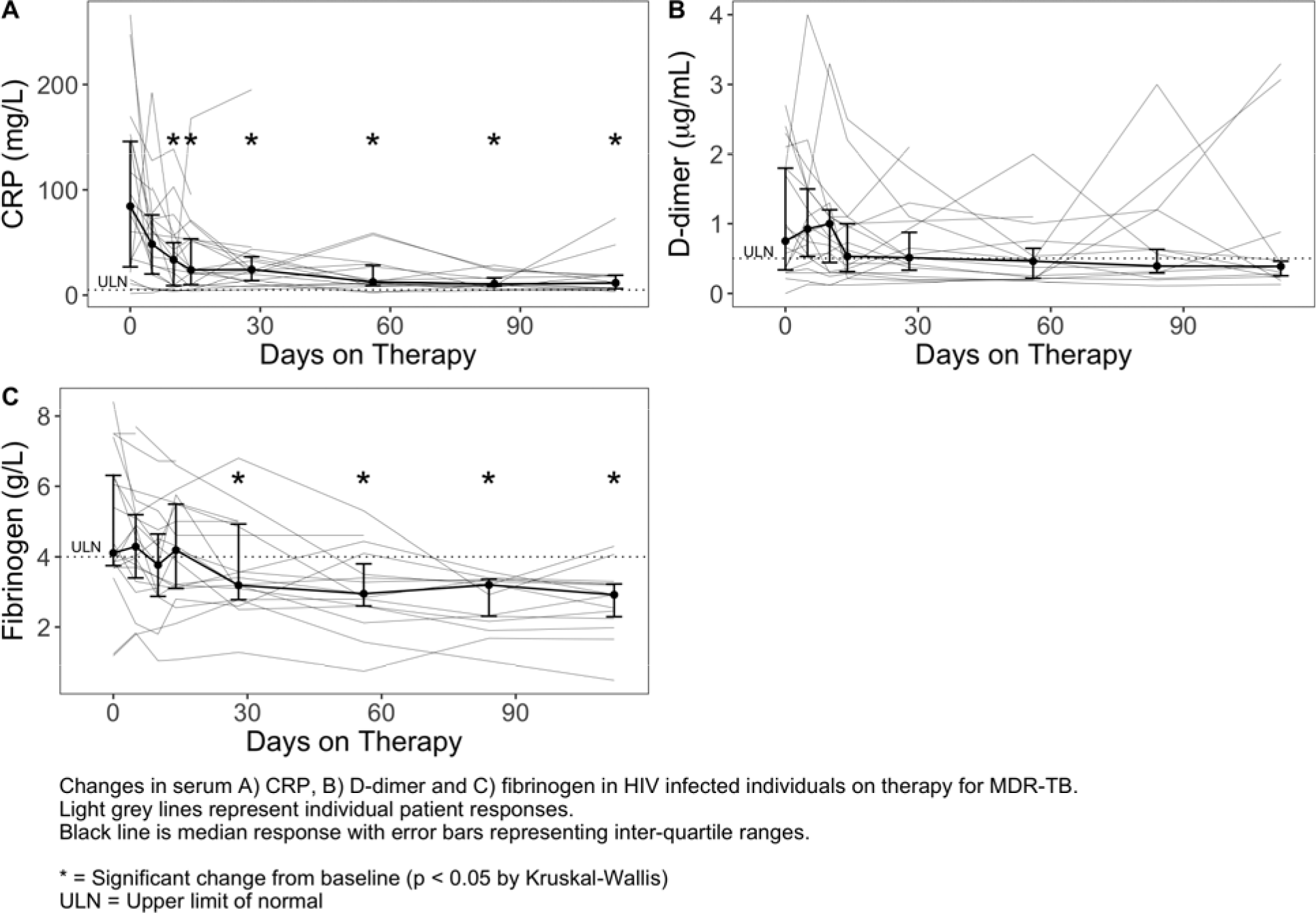
Biomarker trends while on MDR-TB therapy.

